# CD133^+^ progenitor cells promote pulmonary hypertension through CXCR4 signaling

**DOI:** 10.64898/2026.07.29.741640

**Authors:** Zhiqing Wang, Dan Yi, Xinyi Zhang, Jingbo Dai, Xianming Zhang, You-Yang Zhao, Zhiyu Dai

## Abstract

**Background:** Pulmonary hypertension (PH) is characterized by pulmonary vascular remodeling and smooth muscle cell accumulation, but the progenitor-like cells that contribute to this process remain incompletely defined.

**Methods:** We combined analyses of human pulmonary arterial hypertension lungs and experimental PH models with bulk and single-cell RNA sequencing, lineage tracing, inducible ablation of CD133^+^ cells, and conditional deletion of Cxcr4 in CD133^+^ cells.

**Results:** CD133 expression was markedly increased in human and experimental PH lungs. Transcriptomic analyses identified inflammatory, metabolic, chemokine-associated, and smooth muscle cell-like programs in CD133^+^ cells from PH lungs. Lineage tracing showed that CD133^+^ cells contributed to endothelial and smooth muscle cell compartments during experimental PH. Genetic ablation of CD133^+^ cells attenuated hypoxia-induced PH and pulmonary vascular remodeling, whereas Cxcr4 deletion in CD133^+^ cells reduced PH severity.

**Conclusions:** CD133^+^ progenitor cells are functional contributors to pulmonary vascular remodeling, and CXCR4 signaling mediates their pathogenic activity. Targeting pathogenic CD133^+^ cell states or CXCL12/CXCR4 signaling may provide a strategy to limit vascular remodeling in PH.

## Introduction

Pulmonary hypertension (PH) is a progressive cardiopulmonary disorder characterized by pulmonary vascular remodeling, increased pulmonary vascular resistance, right ventricular hypertrophy, and eventual right heart failure. (1) Remodeling of distal pulmonary arteries, including neomuscularization and abnormal accumulation of smooth muscle cells (SMCs), is a central pathologic feature that drives disease progression. (1) Despite substantial progress in defining signaling pathways involved in PH, the cellular origins and molecular mechanisms underlying pathologic SMC expansion remain incompletely understood. (1, 2)

Emerging evidence suggests that progenitor-like cell populations contribute to pulmonary vascular remodeling. (3, 4) Prominin-1 (Prom1), also known as CD133, is a well-established marker of stem and progenitor cells. (5) CD133 was initially identified in a subset of CD34^+^ hematopoietic progenitors from fetal liver, bone marrow, and peripheral blood, including circulating endothelial precursor cells. (6–8) Subsequent studies showed that CD133^+^ stem-like cells are multipotent and can differentiate into multiple lineages. (9, 10) In addition to their differentiation potential, CD133^+^cells can influence disease progression through paracrine signaling. For example, Jagged-1 derived from CD133^+^ endothelial cells activates Notch signaling in colorectal cancer cells and promotes tumor growth. (11) In the lung, bone marrow-derived CD133^+^ endothelial progenitor-like cells have been reported to infiltrate pulmonary arteries in chronic obstructive pulmonary disease and to differentiate into endothelial and smooth muscle cell lineages, suggesting that CD133^+^ cells may participate in vascular remodeling. (3)

Several observations further support a potential role for CD133^+^ cells in PH. CD34-positive/CD133-positive progenitor cells are increased in the bone marrow, peripheral blood, and pulmonary arteries of patients with pulmonary arterial hypertension (PAH) relative to controls. (12) Other studies have also reported elevated CD133 expression and increased numbers of circulating CD133^+^ cells in PAH. (4) However, whether CD133^+^ cells directly contribute to pulmonary vascular remodeling in PH, which CD133^+^ cell states expand in disease, and what signaling pathways regulate their pathogenic activity remain unclear.

In the present study, we examined the role of CD133^+^ cells in PH using human lung tissues together with complementary mouse models, including *Egln1^Tie2Cre^* mice (13) and chronic hypoxia-induced PH. We found that CD133 expression is markedly upregulated in PH and that CD133^+^ cells expand under disease conditions. Integrating bulk RNA sequencing, single-cell RNA sequencing, and lineage-tracing approaches, we identified a marked enrichment of CD133^+^ SMCs and obtained evidence that CD133^+^ progenitor cells contribute to smooth muscle cell accumulation during hypoxic remodeling. Functional studies further showed that depletion of CD133^+^ cells attenuates hypoxia-induced PH, whereas selective deletion of Cxcr4 in CD133^+^ cells protects against disease progression. Together, these data support a role for CD133^+^ progenitor-like cells in pulmonary vascular remodeling and identify CXCR4 signaling as a mediator of their pathogenic function in PH.

## Results

### CD133 expression is increased in human and experimental PH

To determine whether CD133 is associated with PH, we first examined lung tissues from patients with idiopathic PAH (IPAH). PROM1 (encoding CD133) mRNA expression was significantly increased in IPAH lungs relative to control lungs (**Figure 1A**). Consistent with this finding, Prom1/CD133 expression was also elevated in *Egln1^Tie2Cre^* mice, a spontaneous severe PH model, (13) and in Sugen5416/hypoxia-treated mice compared with their respective controls (**Figure 1B, C**). Immunostaining further showed prominent CD133 expression within vascular lesions and muscularized pulmonary arterioles of SuHx mice where CD133 colocalized with α-SMA (**Figure 1D, E**). Immunostaining and flow cytometric analysis likewise demonstrated a markedly increased proportion (33-fold) of CD133^+^/α-SMA^+^ cells in the lungs of *Egln1^Tie2Cre^* mice compared with WT mice (**Figure 1F and G**). Together, these findings show that CD133 expression is increased in both human PAH and experimental PH and is associated with smooth muscle cell-enriched vascular lesions.

**Figure 1.**
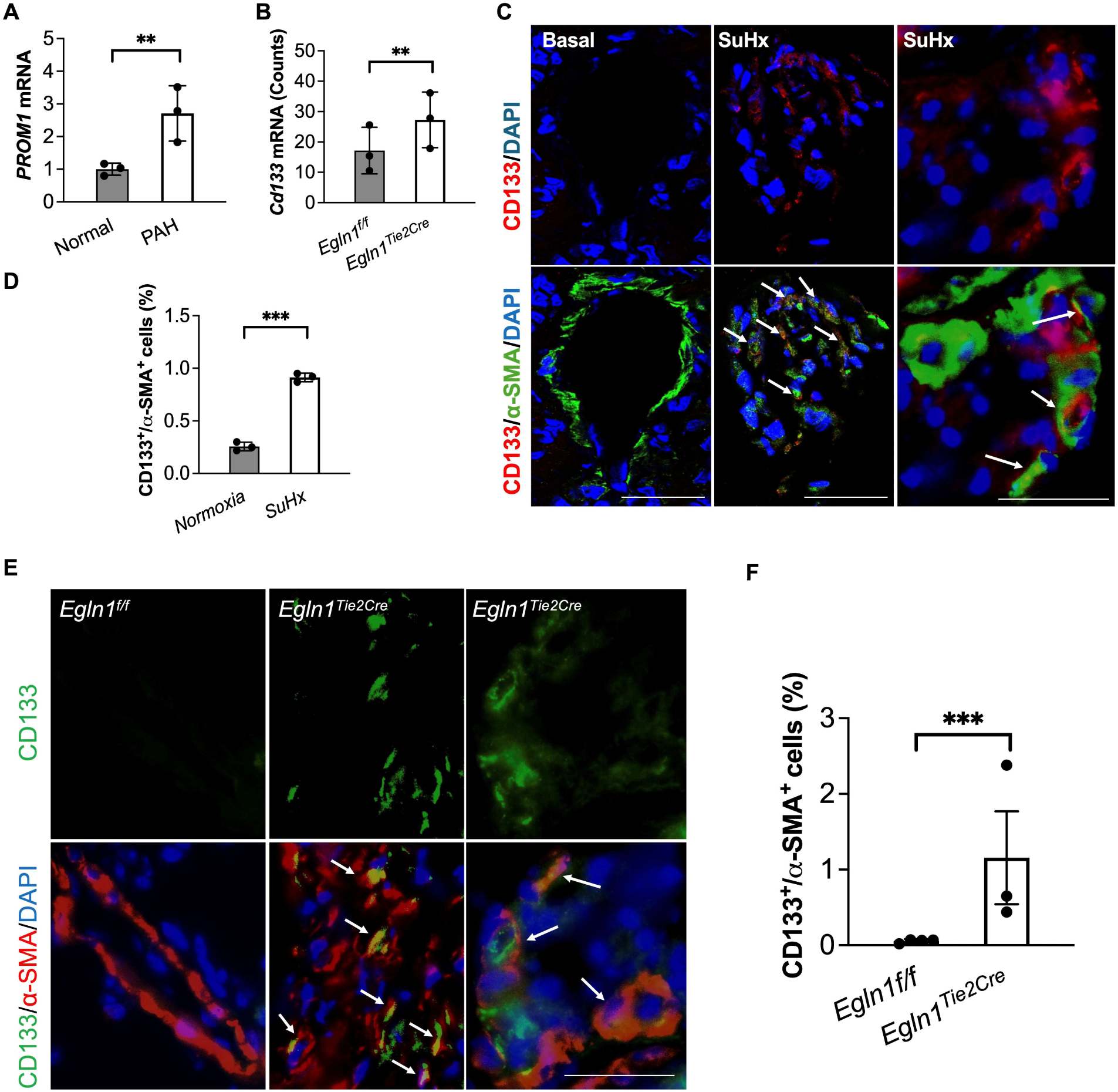
CD133 is upregulated in human and experimental PH mouse models and marks smooth muscle-associated vascular lesions. (A–C) Quantitative RT-PCR analysis of CD133/Prom1 mRNA levels in lung tissues from patients with idiopathic pulmonary arterial hypertension (PAH) and control subjects **(A)**, *Egln1^Tie2Cre^* mice and littermate controls **(B)**. **(C, D)** Representative immunofluorescence images and quantification showing CD133 expression together with α-SMA in pulmonary vascular lesions from Sugen5416/hypoxia-treated mice. (**E**) Representative immunofluorescence images showing CD133 expression together with α-SMA in pulmonary vascular lesions from *Egln1^Tie2Cre^* mice. (**F**) Flow cytometric analysis of CD133^+^/α-SMA^+^ cells in lungs of *Egln1^Tie2Cre^* mice. Scale bars, 50 µm. **P < 0.01; ***P < 0.001. t test (A, B, D, F)

### Transcriptomic profiling of CD133^+^ cells in PH

To define the molecular features of CD133^+^ cells during PH, we isolated CD133^+^ cells using antibody-conjugated microbeads from *Egln1^f/f^* control and *Egln1^Tie2Cre^*lungs and performed bulk RNA sequencing (**Figure 2A, B**). Gene set enrichment analysis revealed that CD133^+^ cells from *Egln1^Tie2Cre^*mice were enriched for inflammatory and stress-response pathways, including interferon-α response, interferon-γ response, TNFα signaling via NF-κB, inflammatory response, and IL6-JAK-STAT3 signaling (**Figure 2C, D**). Metabolic and growth-associated pathways, including oxidative phosphorylation, MYC targets, mTORC1 signaling, and hypoxia-related programs, were also enriched in the *Egln1^Tie2Cre^* group. In contrast, estrogen response early, KRAS signaling DN (down-regulated), and myogenesis were negatively enriched.

**Figure 2.**
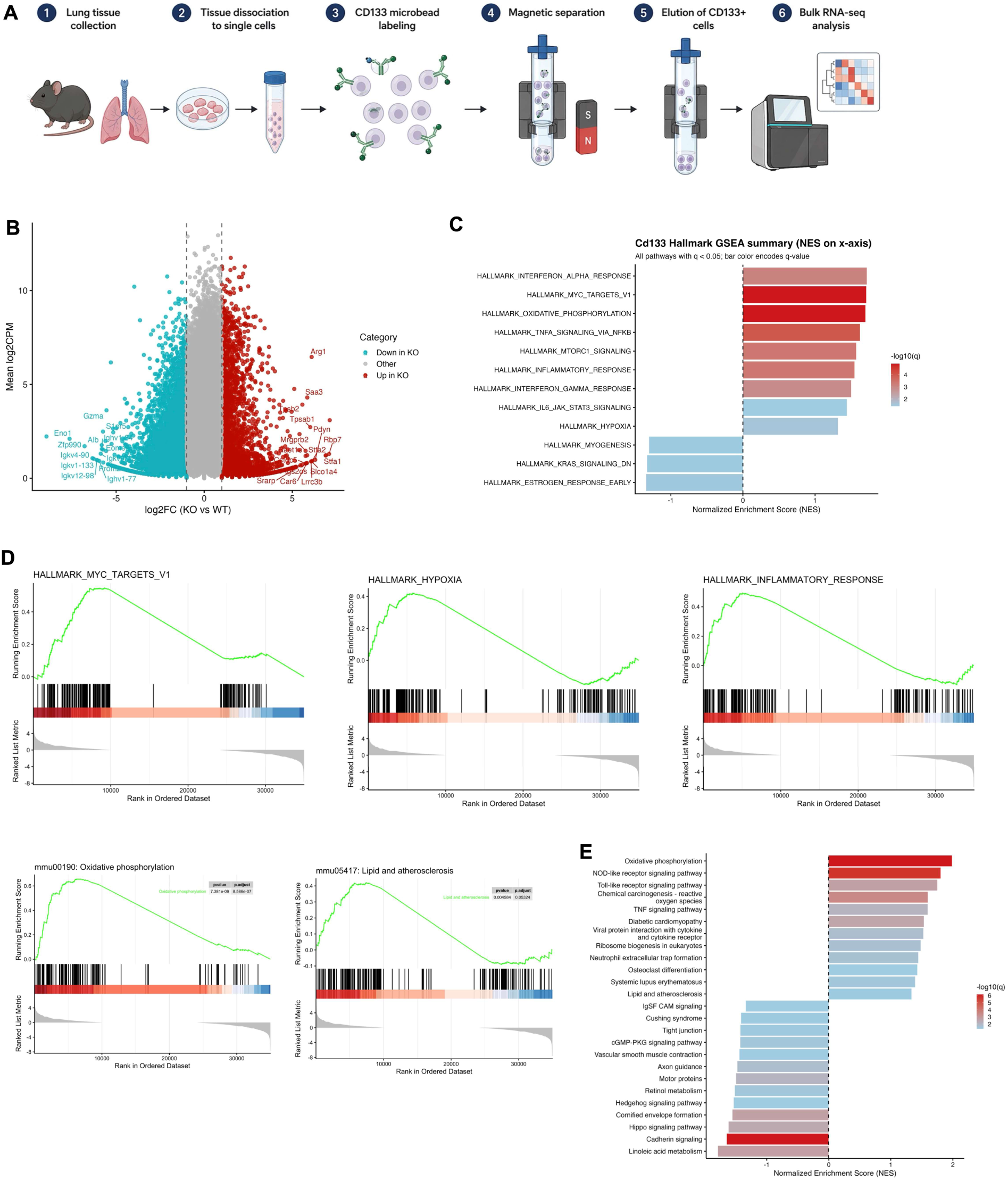
Bulk RNA sequencing reveals inflammatory, metabolic, and CXCL12-associated programs in CD133^+^ cells from. *Egln1*^Tie2Cre^ **PH lungs. (A)** Experimental scheme for isolation of CD133^+^ cells from *Egln1*^f/f^ control and *Egln1*^Tie2Cre^ lungs for bulk RNA sequencing. **(B)** Volcano plot showing differentially expressed genes in CD133^+^ cells from *Egln1*^Tie2Cre^ versus control mice. **(C, D)** Gene set enrichment analysis showing pathways enriched **in** CD133^+^ cells from *Egln1*^Tie2Cre^ mice relative to controls. **(E)** KEGG pathway analysis of differentially regulated pathways in CD133^+^ cells from *Egln1*^Tie2Cre^ mice.

KEGG pathway analysis further supported these findings. Among positively enriched pathways, oxidative phosphorylation, NOD-like receptor signaling, Toll-like receptor signaling, chemical carcinogenesis-reactive oxygen species, and lipid and atherosclerosis pathways were prominent in CD133^+^ cells from *Egln1^Tie2Cre^* mice (**Figure 2E**). By contrast, linoleic acid metabolism, cadherin signaling, Hippo signaling, and retinol metabolism were reduced. These data suggest that CD133^+^ cells in PH acquire an activated transcriptional state characterized by inflammatory signaling, metabolic rewiring, and stress-response pathway engagement.

Heatmap analysis of representative lineage markers showed broadly similar expression of many structural and immune cell markers between groups, but several myeloid/macrophage-associated genes were increased in CD133^+^ cells from *Egln1^Tie2Cre^* mice. Notably, Klf4, a marker linked to smooth muscle cell differentiation, was upregulated in the *Egln1^Tie2Cre^*group, consistent with a disease-associated smooth muscle cell-like program (**Supplemental Figure 1A**). In addition, expression of multiple chemokines, including Ccl5, Ccl6, Cxcl1, and Cxcl2 as well as Cxcl12 was markedly elevated in CD133^+^ cells from *Egln1^Tie2Cre^* mice (**Supplemental Figure 1B**). Given the known role of CXCL12 signaling in vascular remodeling, (14) these data raise the possibility that CD133^+^ cells contribute to PH, at least in part, through CXCL12-associated signaling.

### Single-cell RNA sequencing and lineage tracing of CD133^+^ cells in PH

We next performed single-cell RNA sequencing to resolve the cellular heterogeneity of the isolated CD133^+^ compartment in control and *Egln1^Tie2Cre^* lungs (**Figure 3A**). This analysis showed that CD133^+^ cells represent a heterogeneous population composed of multiple cell states, including endothelial cells, smooth muscle cells, fibroblasts, and alveolar macrophages (**Figure 3B**). Quantitative comparison revealed a marked increase in the proportion of CD133^+^ smooth muscle cells in *Egln1^Tie2Cre^* mice, consistent with expansion of a smooth muscle cell-associated CD133^+^ population during PH (**Supplemental Figure 2A, and Figure 3B, C**).

**Figure 3.**
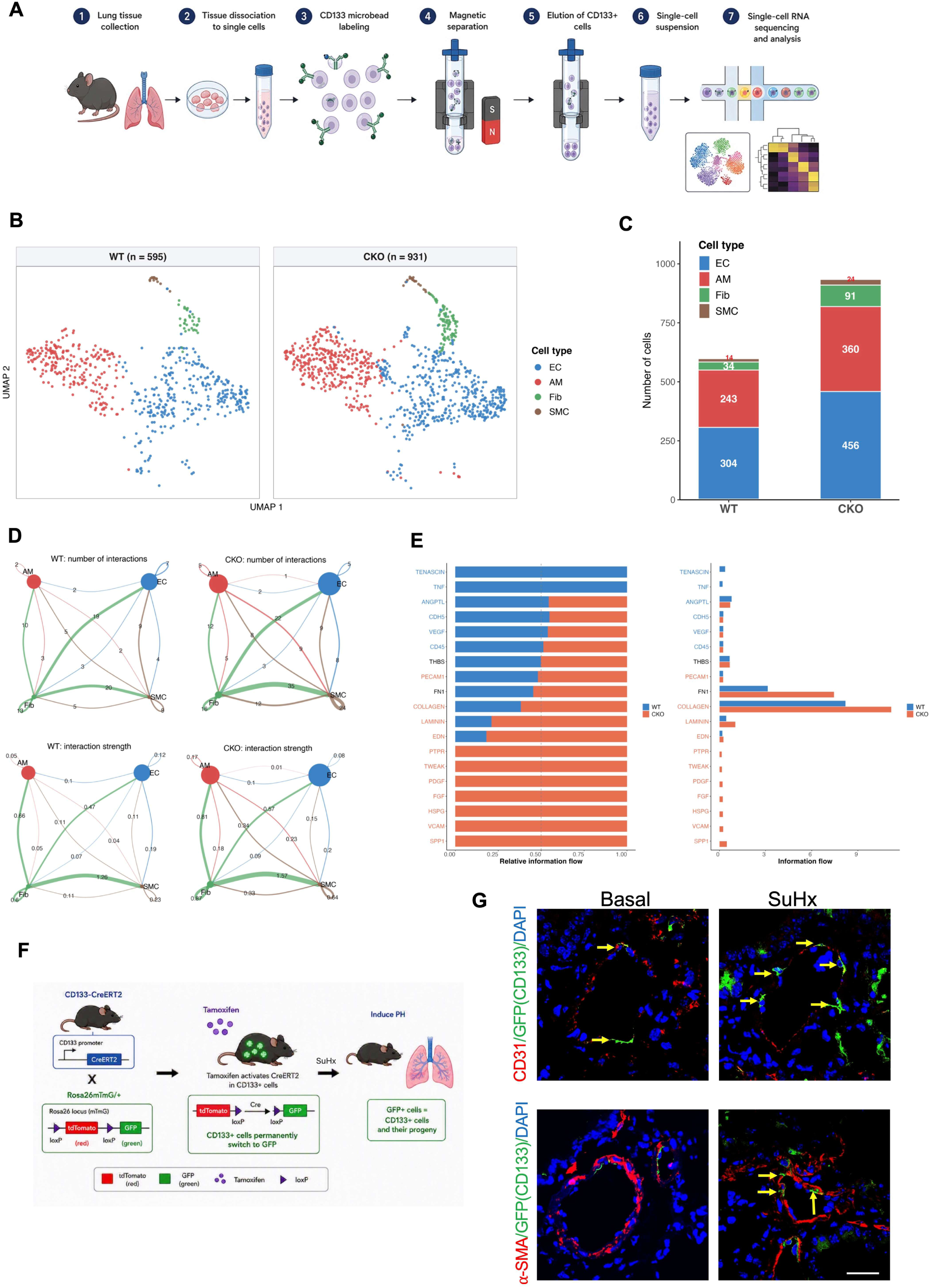
Single-cell RNA sequencing and lineage tracing show expansion and smooth muscle contribution of CD133^+^ cells in PH. (**A**) Graphic presentation of procedures for single-cell RNA sequencing. **(B)** UMAP visualization of CD133^+^ cells isolated from control and *Egln1*^Tie2Cre^ lungs, with subclustering resolving endothelial cell, smooth muscle cell, fibroblast, and alveolar macrophage populations within the CD133^+^ compartment. **(C)** Quantification of major CD133^+^ cell populations in control and *Egln1*^Tie2Cre^ mice. **(D)** Predicted cell-cell communication networks among CD133^+^ cell populations in control and *Egln1*^Tie2Cre^ lungs. (**E**) Ligand-receptor interaction among CD133^+^ cell populations in control and *Egln1*^Tie2Cre^ lungs. **(F)** Schematic of tamoxifen-inducible lineage tracing in *Cd133*-CreERT2;*Rosa26*mTmG/+ mice. SuHx, Sugen5416/hypoxia. **(G)** Representative immunofluorescence images showing lineage-labeled CD133^+^ cells within CD31^+^ endothelial and α-SMA^+^ smooth muscle compartments. Scale bar, 50 μm (**G**)

We then examined predicted cell-cell communication among CD133^+^ cell states using CellChat. (24) Compared with WT lungs, *Egln1^Tie2Cre^* lungs exhibited more predicted interactions (176 versus 113) and greater overall interaction strength (6.34 versus 4.15). Pathway-level analysis showed increased information flow through COLLAGEN, LAMININ, and FN1 signaling, with SPP1, FGF, and PDGF signaling detected only in the *Egln1^Tie2Cre^* group. In a CXCL-focused sensitivity analysis, CXCL signaling communication was detected only in *Egln1^Tie2Cre^* lungs, with endothelial cells, macrophages, fibroblasts, and smooth muscle cells predicted as CXCL sources. (**Figure 3D and Supplemental Figure 2B**). CXCL signaling emerged as a prominent component of this altered communication network, further supporting a role for chemokine signaling in CD133^+^ cell-associated remodeling (**Figure 3E**).

To determine the fate of CD133^+^ cells during PH, we performed lineage tracing using tamoxifen-inducible *Cd133*-CreERT2;*Rosa26*mTmG/+ reporter mice (15) (**Figure 3F, G**). After Sugen5416/hypoxia exposure, lineage-labeled CD133^+^ cells were detected in both CD31^+^ endothelial and α-SMA^+^ smooth muscle compartments (**Figure 3G**). These findings indicate that CD133^+^ cells expand during PH and support the interpretation that CD133^+^ progenitor cells contribute to endothelial and smooth muscle cell accumulation during vascular remodeling.

### Depletion of CD133^+^ cells attenuates hypoxia-induced PH and remodeling

To test whether CD133^+^ cells functionally contribute to PH, we generated inducible CD133^+^ cell ablation mice by crossing *Cd133*-CreERT2 mice with *Rosa26*iDTR/+ mice and then induced diphtheria toxin receptor expression with tamoxifen. (16) Diphtheria toxin treatment was then used to ablate CD133^+^ cells (**Figure 4A**). Flow cytometry showed that approximately 90% of CD133^+^ cells were depleted within 3 days, and this reduction persisted for at least 10 days (**Figure 4B**). Following chronic hypoxia exposure, mice with CD133^+^ cell depletion exhibited significantly lower right ventricular systolic pressure (RVSP) and reduced right ventricular hypertrophy compared with control mice (**Figure 4C, D**).

**Figure 4.**
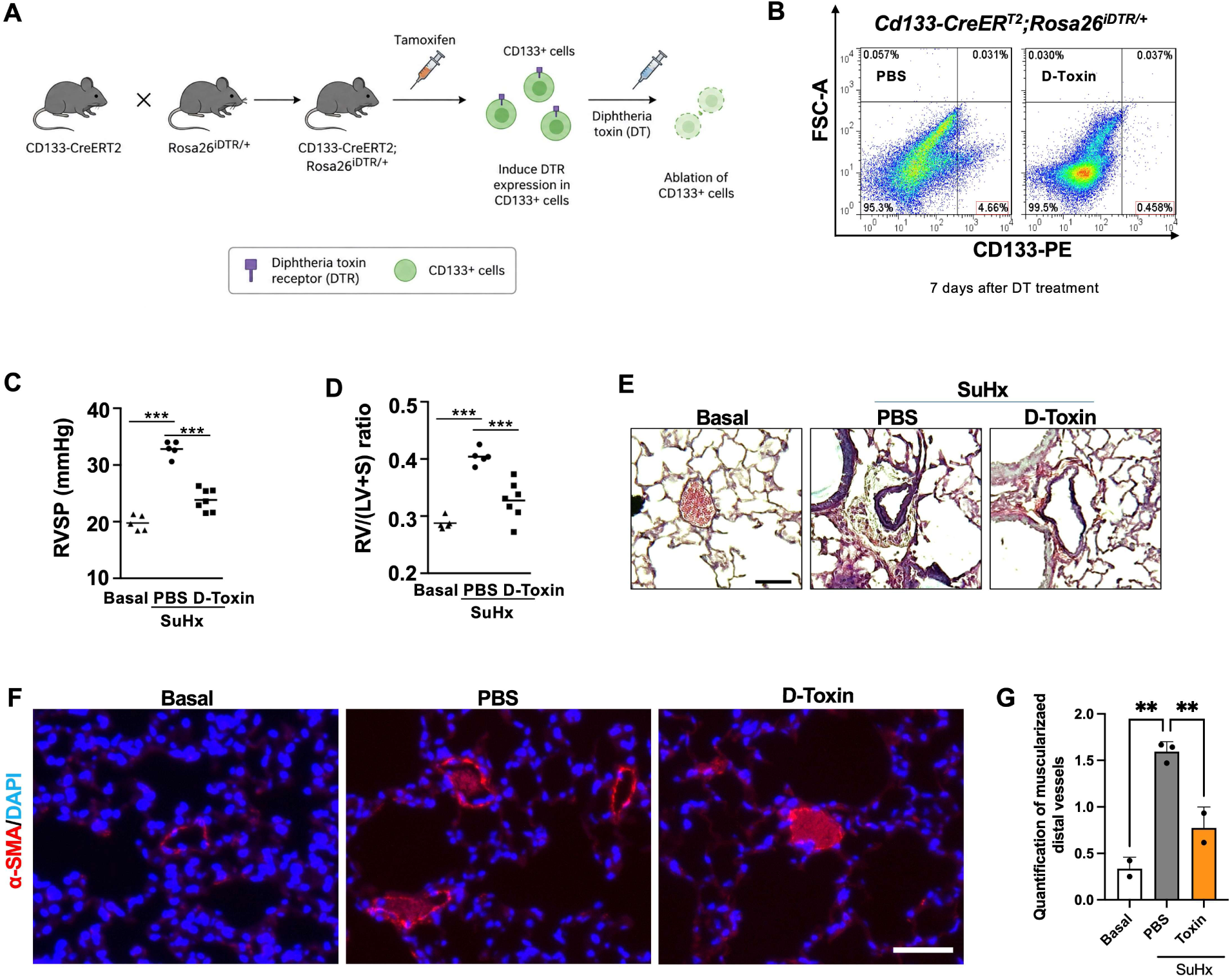
Ablation of CD133^+^ cells attenuates hypoxia-induced pulmonary vascular remodeling and PH. **(A)** Schematic diagram of diphtheria toxin-mediated ablation of CD133^+^ cells in *Cd133*-CreERT2;*Rosa26*iDTR/+ mice. **(B)** Flow cytometric analysis of CD133^+^ cell depletion after diphtheria toxin treatment. **(C)** Right ventricular systolic pressure (RVSP). **(D)** Right ventricular hypertrophy, expressed as RV/(LV+S) ratio. **(E)** Representative Russell-Movat pentachrome staining of lung sections. **(F, G)** Representative micrographs of α-SMA staining and quantification of muscularized distal vessels (diameters ≤50 µm) and quantification of SMA^+^ full coverage vessels. ***P < 0.001. One-way ANOVA with Tukey post-hoc analysis (**C**, **D, G**). Scale bar, 50 μm (**E**, **F**).

Histologic analysis further showed that pulmonary arterial wall thickening was reduced after CD133^+^ cell depletion (**Figure 4E**). Consistent with this finding, α-SMA staining demonstrated decreased smooth muscle coverage in distal small (<50 µm) vessels in mice lacking CD133^+^ cells (**Figure 4F–G**). Together, these data indicate that CD133^+^ cells contribute functionally to hypoxia-induced pulmonary vascular remodeling and PH.

### Cxcr4 deletion in CD133^+^ cells reduces hypoxia-induced PH

Because Cxcl12 expression was increased in CD133^+^ cells and CXCL12 signaling has been implicated in pulmonary vascular remodeling, (14) we next tested whether Cxcr4 deletion in CD133^+^ cells alters PH severity. *Cd133*-CreERT2 mice were crossed with *Cxcr4*^f/f^ mice to generate *Cd133*-CreERT2;*Cxcr4*^f/f^ mice (**Figure 5A**). Compared with littermate controls, mice lacking Cxcr4 in CD133^+^ cells exhibited significantly lower RVSP and reduced right ventricular hypertrophy after hypoxia exposure (**Figure 5B-D**). These findings support a role for CXCL12-CXCR4 signaling in CD133^+^ cell-mediated PH and suggest that this pathway contributes to the pathogenic activity of CD133^+^ cells during vascular remodeling.

**Figure 5.**
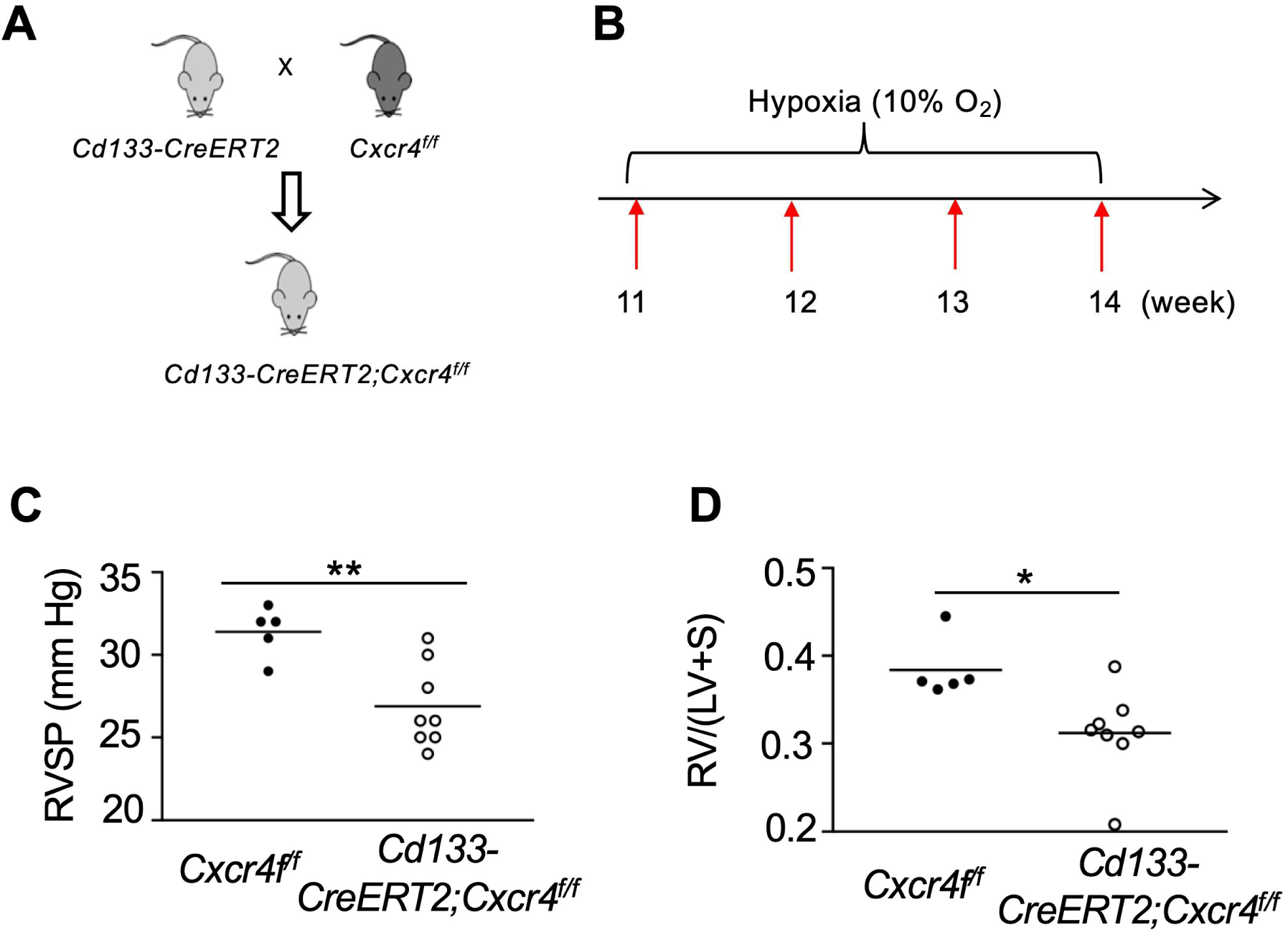
Cxcr4 deletion in CD133^+^ cells protects against hypoxia-induced PH. **(A)** Breeding strategy for generation of *Cd133*-CreERT2;*Cxcr4*^f/f^ mice. (**B**) Experimental design for hypoxia exposure of *Cd133*-CreERT2;*Cxcr4*^f/f^ mice and littermate controls. **(C)** Right ventricular systolic pressure (RVSP). **(D)** Right ventricular hypertrophy, expressed as RV/(LV+S) ratio. *P < 0.05; **P < 0.01. t test (**C**, **D**).

## Discussion

Pulmonary vascular remodeling in PH is characterized by progressive muscularization of distal pulmonary arteries, accumulation of smooth muscle-like cells, inflammatory and metabolic activation of the vascular wall, and increased pulmonary vascular resistance that ultimately drives right heart failure. (1) Although expansion of pre-existing vascular smooth muscle cells is an important component of this process, several studies have suggested that additional progenitor-like populations also participate in neomuscularization. (17–19) In the present study, we identify CD133^+^ cells as a disease-associated progenitor-like population that expands during PH and contributes functionally to vascular remodeling. CD133 expression was increased in human PAH lungs and in complementary mouse PH models; CD133^+^ cells acquired inflammatory, metabolic, and chemokine-associated transcriptional programs; CD133^+^ smooth muscle-like cells were expanded in diseased lungs; and genetic depletion of CD133^+^ cells attenuated hypoxia-induced PH. Together, these findings move CD133 beyond a disease-associated marker and support a model in which CD133^+^ progenitor-like cells participate directly in pulmonary vascular remodeling.

These data extend prior observations linking CD133^+^ cells to pulmonary vascular disease. CD133 has long been used to identify stem and progenitor populations in hematopoietic and endothelial compartments, (5–8) and CD133^+^ cells can display substantial lineage plasticity and paracrine activity in tissue remodeling and cancer. (9–11) In the lung, CD133^+^ bone marrow-derived cells have been reported to infiltrate remodeled pulmonary arteries in chronic obstructive pulmonary disease and to acquire endothelial and smooth muscle features, (3) whereas CD34^+^CD133^+^ progenitor-like cells have been detected in endarterectomized tissue from chronic thromboembolic PH. (20) In PAH, increased CD34^+^CD133^+^ progenitor cells have been reported in bone marrow, peripheral blood, and pulmonary arteries, (12) and elevated circulating CD133^+^ cells have been associated with PAH. (4) Experimental hypoxia has also been shown to increase CD133^+^ progenitor-like cells in pulmonary hypertensive rat lungs, with evidence that these cells can differentiate toward smooth muscle-like cells. (21) However, most of these studies were descriptive, focused on circulating progenitors, or did not directly test whether CD133^+^ cells are required for remodeling. Our lineage tracing and ablation studies address this gap by showing that CD133^+^ cells contribute to the endothelial and smooth muscle compartment and that loss of CD133^+^ cells reduces hemodynamic and structural features of hypoxia-induced PH.

The finding that CD133^+^ cells contribute to smooth muscle cell accumulation is consistent with a broader shift in the field toward multiple cellular origins of distal muscularization. Sheikh and colleagues identified primed smooth muscle progenitors at the muscular–unmuscular border of pulmonary arterioles that migrate, dedifferentiate, clonally expand, and generate distal smooth muscle cells during hypoxia-induced PH. (17) Dierick and colleagues identified resident PW1^+^ progenitor cells that are recruited during early hypoxic remodeling, differentiate into vascular smooth muscle cells, and are increased in remodeled vessels from patients with PAH. (18) Subsequent work further implicated PDGFRα signaling in PW1^+^ progenitor proliferation and vascular remodeling. (22) Our data place CD133^+^ cells within this evolving framework of progenitor-associated neomuscularization. Importantly, the CD133^+^ compartment was heterogeneous by single-cell RNA sequencing, including endothelial, smooth muscle, fibroblast, and macrophage-associated states. Thus, CD133 should not be interpreted as marking a single lineage-restricted precursor. Rather, in PH, CD133 appears to identify a remodeling-associated cellular state that includes a smooth muscle-biased subpopulation capable of contributing to vascular muscularization.

The transcriptomic profile of CD133^+^ cells provides clues to how this population may become pathogenic. CD133^+^ cells from *Egln1^Tie2Cre^* lungs were enriched for interferon responses, TNFα/NF-κB signaling, inflammatory response, IL6-JAK-STAT3 signaling, oxidative phosphorylation, MYC targets, mTORC1 signaling, and hypoxia-associated programs. These pathways align with major themes in PAH pathobiology, in which inflammation, altered metabolism, proliferative signaling, and hypoxia-responsive programs interact to sustain vascular remodeling. (1, 2, 13) The upregulation of Klf4 is also notable because KLF4 has been implicated in smooth muscle phenotypic modulation and in primed smooth muscle progenitor behavior during hypoxic muscularization. (17) In this context, the expansion of CD133^+^ smooth muscle-like cells in our single-cell dataset suggests that CD133^+^ cells are not only more abundant in PH but are transcriptionally remodeled toward a state compatible with inflammatory activation, survival, proliferation, and smooth muscle accumulation.

A central mechanistic finding of this study is the involvement of CXCL12/CXCR4 signaling in CD133^+^ cell-mediated PH. We found increased Cxcl12 expression in CD133^+^ cells from *Egln1^Tie2Cre^* lungs, prominent CXCL signaling in predicted cell-cell communication networks, and protection from hypoxia-induced PH after Cxcr4 deletion in CD133^+^ cells. These findings agree with prior evidence that CXCL12/CXCR4 signaling contributes to pulmonary vascular remodeling. (14) They also resonate with the PW1^+^ progenitor study, in which pharmacologic CXCR4 inhibition with AMD3100 prevented differentiation of PW1^+^ progenitor cells into smooth muscle cells during hypoxic remodeling. (18) Our genetic approach extends these observations by assigning functional importance to CXCR4 within the CD133^+^ compartment itself. Although the current study does not determine whether CXCL12 acts through autocrine signaling among CD133^+^ cells, paracrine signaling from neighboring vascular cells, or both, the protection observed after CD133-restricted Cxcr4 deletion supports CXCR4 as a regulator of the pathogenic activity of this population.

The functional studies strengthen the biological significance of the expression and transcriptomic findings. Ablation of CD133^+^ cells reduced RVSP, right ventricular hypertrophy, pulmonary arterial wall thickening, and α-SMA^+^ muscularization in hypoxia-exposed mice. These results indicate that CD133^+^ cells are not merely a correlation of disease severity but contribute to PH pathogenesis. At the same time, the data should be interpreted with appropriate caution. CD133 is expressed by multiple progenitor-like and differentiated cell states, (3, 5) and the protection observed after cell depletion likely reflects the combined loss of several CD133^+^ subpopulations. In addition, CD133 expression may mark an activated state rather than define a fixed cell lineage. Nevertheless, the convergence of human tissue analysis, mouse PH models, single-cell profiling, lineage tracing, cell ablation, and CD133-restricted Cxcr4 deletion provides complementary evidence that CD133^+^ progenitor-like cells are functionally relevant contributors to pulmonary vascular remodeling.

This study has several limitations. First, although lineage tracing supports contribution of CD133^+^ cells to the smooth muscle compartment, the relative contribution of resident vascular CD133^+^ cells versus recruited circulating progenitor-like cells remains unresolved. Second, the molecular mechanisms that connect CD133 expression, CXCR4 signaling, KLF4-associated programs, and smooth muscle differentiation require further study. Third, the ablation and conditional knockout experiments were performed in hypoxia-based PH, and additional work will be needed to determine whether the same mechanisms operate in severe angioproliferative or obliterative models, including *Egln1^Tie2Cre^* mice and Sugen5416/hypoxia rats. Fourth, because CD133^+^ cells are heterogeneous, future studies using higher-resolution lineage tracing, spatial transcriptomics, and subpopulation-specific genetic tools will be important to identify the specific CD133^+^ states that are necessary and sufficient for remodeling.

In summary, our findings identify CD133^+^ cells as a disease-associated progenitor-like population that expands in PH, acquires inflammatory and remodeling-associated transcriptional programs, contributes to smooth muscle accumulation, and promotes pulmonary vascular remodeling through CXCR4-dependent mechanisms. These results refine current understanding of the cellular origins of vascular muscularization in PH and suggest that targeting pathogenic CD133^+^ cell states or their CXCL12/CXCR4 signaling programs may represent a strategy for limiting pulmonary vascular remodeling.

## Methods

### Human Samples

The use of archived human lung tissues was granted by the institutional review board. Human patients with idiopathic PAH and healthy donors were obtained from the Pulmonary Hypertension Breakthrough Initiative (PHBI), with informed consent and institutional review board approval at the transplant procurement sites. The clinical and demographic characteristics of patients with IPAH and control is described previously (13).

### Animals and experimental design

All animal studies were performed on a C57BL/6J background using mice maintained on standard chow. *Egln1*^Tie2Cre^ mice were generated as described previously. (13) Male and female littermate controls at 5 weeks to 3.5 months of age were used.

*Cd133-*CreERT2 (JAX #030383), *Rosa26*mTmG/+ (JAX #037456), *Rosa26*iDTR/+ (JAX#008040), and *Cxcr4*^f/f^ (JAX #008767) mice were obtained from The Jackson Laboratory. *Rosa26*mTmG/+ (15), *Rosa26*iDTR/+ (16), and *Cxcr4*^f/f^ mice were crossed with *Cd133-*CreERT2 mice to generate lineage-tracing, ablation, and conditional *Cxcr4*-deletion cohorts, respectively. For lineage-tracing and conditional deletion studies, 7-week-old mice received tamoxifen (100 mg/kg, intraperitoneally) daily for 5 days, followed by a 3-week washout period before exposure to 10% hypoxia for 3 weeks. For CD133^+^ cell-ablation studies, offspring of *Cd133*-CreERT2 crossed with Rosa26iDTR/+ mice received tamoxifen as above, followed by diphtheria toxin (20 ng/g body weight, intraperitoneally) for 3 consecutive days to deplete CD133^+^ cells. (16) All animal procedures were approved by the Institutional Animal Care and Use Committees of Northwestern University and Washington University in St. Louis.

### Hemodynamic measurements and right ventricular hypertrophy determination

RVSP was measured using a 1.4F pressure transducer catheter (Millar Instruments) connected to AcqKnowledge software (Biopac Systems Inc), as described previously. (13) Briefly, under ketamine/xylazine anesthesia (100/5 mg/kg, intraperitoneally), the catheter was inserted into the right ventricle through the jugular vein for pressure recording. After hemodynamic measurements, hearts were harvested and dissected under a microscope to separate the right ventricle (RV) from the left ventricle plus septum (LV+S). Right ventricular hypertrophy was assessed as the RV/(LV+S) weight ratio.

### Immunofluorescence and histology

For immunofluorescence, lungs were perfused with PBS, inflated with 50% optimal cutting temperature compound in PBS, embedded in optimal cutting temperature compound, and cryosectioned at 5 µm. Sections were fixed in 4% paraformaldehyde, blocked with 0.1% Triton X-100 and 5% normal goat serum for 1 hour at room temperature, and incubated overnight at 4^°^C with primary antibodies including anti-α-smooth muscle actin (α-SMA; Abcam, 1:300) and anti-CD133 (Invitrogen, Cat# 46-1331-82). Sections were then incubated with Alexa Fluor 594-, 488-, or 647-conjugated secondary antibodies (Thermo Fisher Scientific) for 1 hour at room temperature and mounted with DAPI-containing medium. Quantification of α-SMA^+^ vessels was performed blindly to genotype and treatment group.

For histology, lungs were perfused with PBS, fixed by tracheal instillation of 10% formalin at a constant pressure of 15 cm H_2_O, embedded in paraffin, and sectioned. Sections were stained with Russell-Movat pentachrome according to the manufacturer’s instructions (StatLab). Pulmonary arterial wall thickness was quantified blindly from 40 images acquired at ×20 magnification using ImageJ. Wall thickness was calculated as the distance between the inner and outer vessel wall divided by the distance from the outer wall to the center of the lumen. Images were acquired using an ECHO REVOLVE microscope (Discover Echo Inc). α-SMA^+^ vessels were quantified in 40 fields per lung.

### Flow cytometry

For flow cytometric analysis, mouse lungs were perfused free of blood with PBS, minced into small pieces, and digested with collagenase A (1 mg/mL; Roche Applied Science) for 1 hour at 37^°^C in a shaking water bath (200 rpm), adapted from the lung single-cell preparation workflow described by Zhang et al. (23) Digested tissues were dissociated using a gentleMACS Dissociator (Miltenyi Biotec), filtered through a 40-µm nylon cell strainer, and blocked with 20% fetal bovine serum for 30 minutes. Cells were then incubated with Fc blocker (1 µg/10^6^ cells; BD Biosciences) for 15 minutes, washed, and resuspended in FACS buffer (PBS containing 5% fetal bovine serum and 5 mM EDTA) and stained for anti-CD133-PE (Miltenyi, Cat# 130-117-784). Red blood cells were lysed with ACK buffer where needed. For depletion studies, flow cytometry was used to quantify the efficiency of CD133^+^ cell ablation after diphtheria toxin treatment. For phenotyping studies, lung cells were stained with antibodies against the indicated surface and lineage markers and analyzed to quantify CD133^+^ and CD133^+^/α-SMA^+^ cell populations.

### Quantitative RT-PCR

Total RNA was isolated from mouse and human tissues using Quick-RNA Miniprep kits with DNase I digestion (Zymo Research). One microgram of RNA was reverse-transcribed using High-Capacity cDNA Reverse Transcription Kits (Applied Biosystems). Quantitative RT-PCR was performed on a QuantStudio 3 system using PowerTrack SYBR Green Master Mix (Applied Biosystems). Relative mRNA expression was calculated by the ΔΔCt method using cyclophilin as the internal control for mouse samples and 18S rRNA for human samples. Primer sequences are listed in Table S1.

### Isolation of CD133-positive cells and bulk RNA sequencing

Lungs were collected from *Egln1*^f/f^ control and *Egln1*^Tie2Cre^ mice after perfusion with HBSS. Tissue was minced and digested with collagenase A (1 mg/mL; Roche) at 37^°^C with shaking for 60 minutes, followed by mechanical dissociation using a Miltenyi tissue dissociator. Consistent with previously described endothelial and vascular tissue dissociation methods, (23) the resulting suspension was filtered through a 40-µm strainer, subjected to red blood cell lysis, and washed in FACS buffer. CD133^+^ cells were isolated using CD133 microbeads and LS columns (Miltenyi Biotec) according to the manufacturer’s protocol.

Total RNA from isolated CD133^+^ cells from 3 control and 3 *Egln1*^Tie2Cre^ mice was extracted using Quick-RNA Miniprep kits with DNase I digestion (Zymo Research). Libraries were sequenced at Novogene on an Illumina NovaSeq 6000 platform with paired-end 150-bp reads. Raw reads were assessed with FastQC, trimmed to remove adapters and low-quality bases (Phred score <33), and aligned to the mouse mm10 genome using STAR (v2.5.2). Gene-level counts were generated with HTSeq using the mm10 Ens.78 annotation. Differentially expressed genes were identified with edgeR (v4.0.1) using an adjusted P <0.05 and log fold change >0.58. Heat maps were generated from scaled RPKM counts using pheatmap (v1.0.12).

### Single-cell RNA sequencing analysis

For single-cell RNA sequencing, CD133^+^ cells from 3 control and 3 *Egln1*^Tie2Cre^ mice were labeled with Cell Hashing antibodies and loaded onto the 10x Genomics Chromium platform for single-cell library preparation. Sequencing was performed using paired-end 150-bp reads, and Cell Ranger (v4.0.0) was used for demultiplexing and count generation against the refdata-gex-mm10 reference genome.

Downstream analyses were performed in Seurat (v4.3.0). Cells with fewer than 200 detected genes or low complexity (log_10_ genes/UMI <0.8) were excluded, and genes expressed in fewer than 3 cells were removed. Doublets were identified using scDblFinder (v1.16.0) and by manual inspection of mixed-lineage marker expression. After doublet removal, datasets were normalized and integrated using an anchor-based canonical correlation analysis pipeline. The first 30 principal components were used for clustering at a resolution of 0.5, and Uniform Manifold Approximation and Projection was used for visualization. Cluster marker genes were identified with FindAllMarkers, and cluster identities were assigned based on canonical marker genes using AddModuleScore. Differential expression between groups within clusters of interest was performed using the Wilcoxon rank-sum test with thresholds of log fold change >0.25, adjusted P <0.05, and minimum cell fraction >0.1.

### Cell-cell communication analysis

Cell-cell communication was inferred from normalized RNA expression matrices using CellChat (v2.1.2) and CellChatDB.mouse. (24) Endothelial cells, alveolar macrophages, fibroblasts, and smooth muscle cells from control and *Egln1^Tie2Cre^* lungs were analyzed separately. All interaction categories in the mouse database, including secreted signaling, extracellular matrix–receptor, cell-cell contact, and nonprotein signaling, were included. Overexpressed signaling genes and ligand-receptor pairs were identified using the standard CellChat workflow. Communication probabilities and permutation P values were calculated with the trimean estimator (100 permutations) using normalized expression data and without weighting by cell population size; interactions involving cell populations with fewer than 10 cells were excluded. Pathway-level probabilities were computed and aggregated, and group-specific CellChat objects were merged to compare the number and strength of interactions and overall signaling information flow.

## Statistical analysis

Statistical analyses were performed using Prism 10 (GraphPad Software). Comparisons among more than 2 groups were performed using one-way ANOVA for normally distributed data, followed by Tukey post hoc testing. Two-group comparisons were analyzed using an unpaired 2-tailed t test. A P value <0.05 was considered statistically significant. Data are presented as mean ± SD.

## Data availability

Bulk RNA sequencing datasets are deposited in NCBI GEO databank (GSE334423). Additional data supporting the findings of this study are available from the corresponding authors upon reasonable request.

## Author contributions

ZW, DY, JD, XZ, YYZ, and ZD contributed to study design, data acquisition, data analysis, interpretation, and manuscript preparation.

## Funding support

This work was supported by National Institutes of Health/National Heart, Lung, and Blood Institute grants R01HL140409, R01HL148810, R01HL133951, R01HL164014, R01HL162299, and R01HL172447 to You-Yang Zhao and R00HL138278, R01HL158596, R01HL162794, R01HL169509, and R01HL170096 to Zhiyu Dai.

## Acknowledgments

None

## Supplemental Figure legends

**Supplemental Figure 1.**
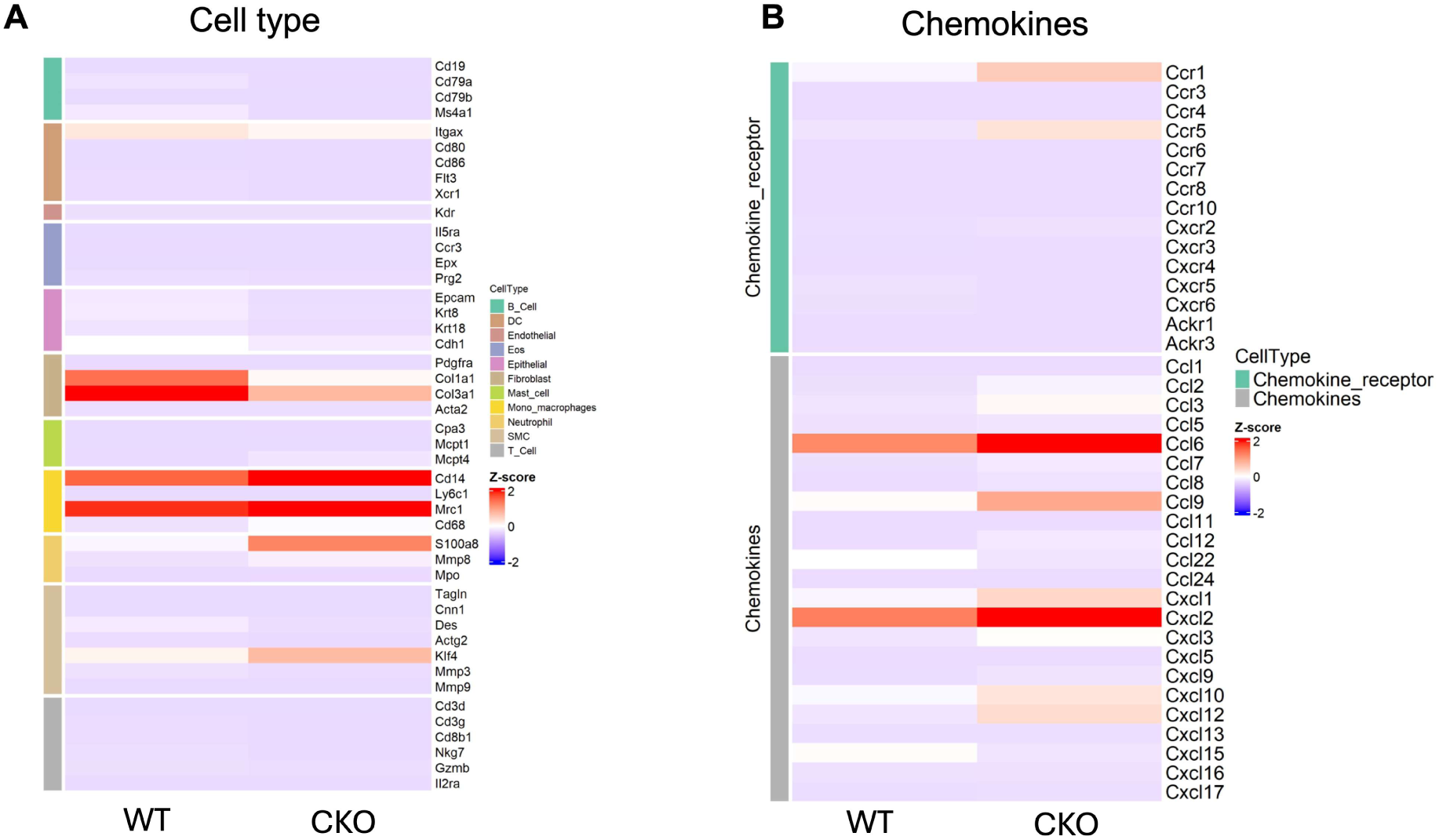
Bulk-RNAseq analysis of CD133^+^ cells. (**A, B**) Representative heatmap for the transcriptomics from CD133^+^ populations.

**Supplemental Figure 2.**
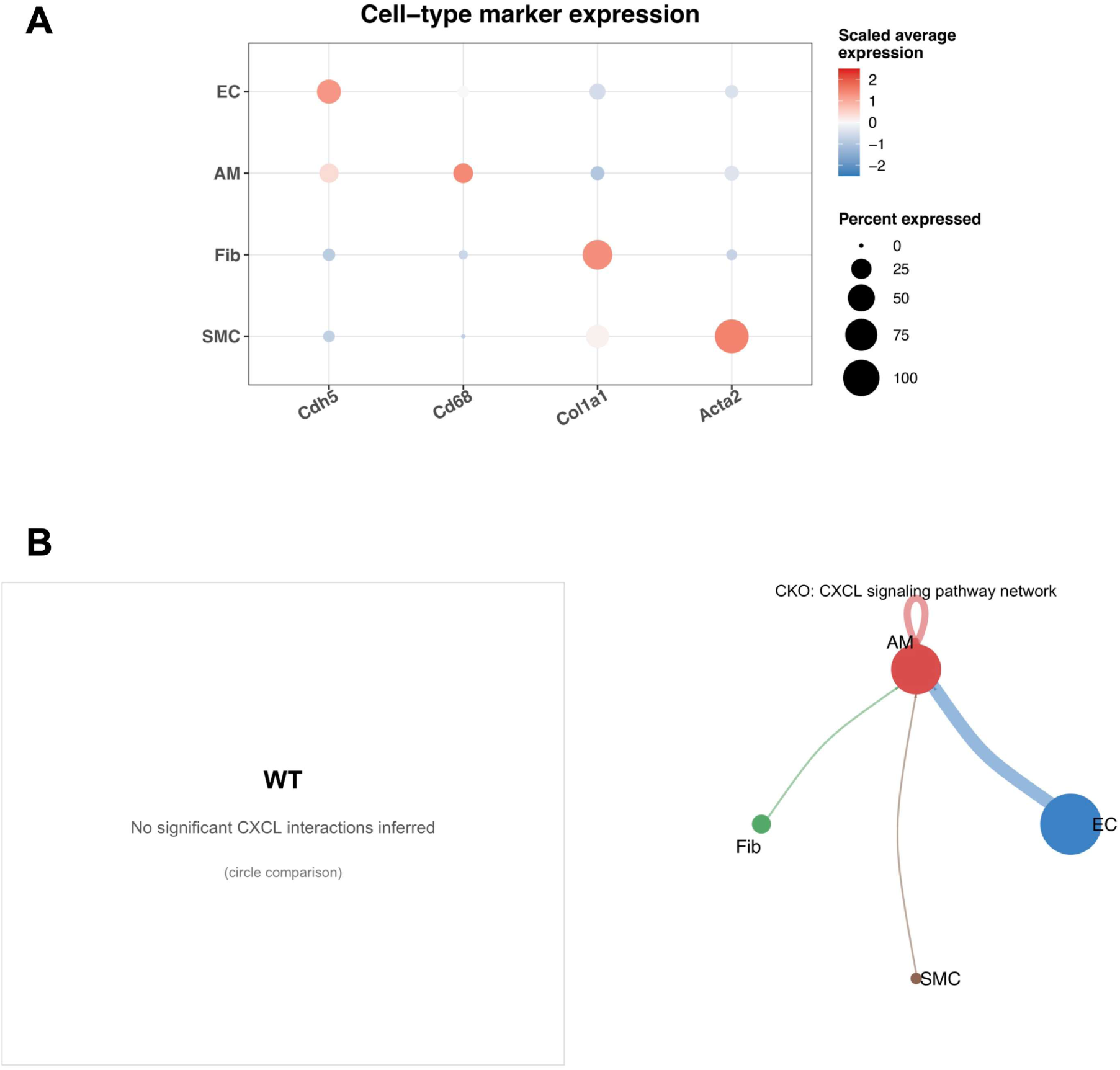
Canonical marker expression and predicted CXCL signaling in CD133^+^ cell populations. (A) Dot plot showing expression of canonical markers for endothelial cells (ECs; Cdh5), alveolar macrophages (AMs; Cd68), fibroblasts (Fib; Col1a1), and smooth muscle cells (SMCs; Acta2). Dot size indicates the percentage of cells expressing each gene, and color represents scaled average expression. (B) CellChat circle plots comparing predicted CXCL signaling between WT and *Egln1^Tie2Cre^* (CKO) CD133^+^ cells. No significant CXCL interactions were inferred in WT cells. In CKO cells, CXCL signaling was predicted from ECs, AMs, fibroblasts, and SMCs toward AMs, including autocrine AM signaling. Node size is proportional to cell number, edge width represents predicted communication probability, and edge color denotes the sending cell population.

---

**Table S1.**
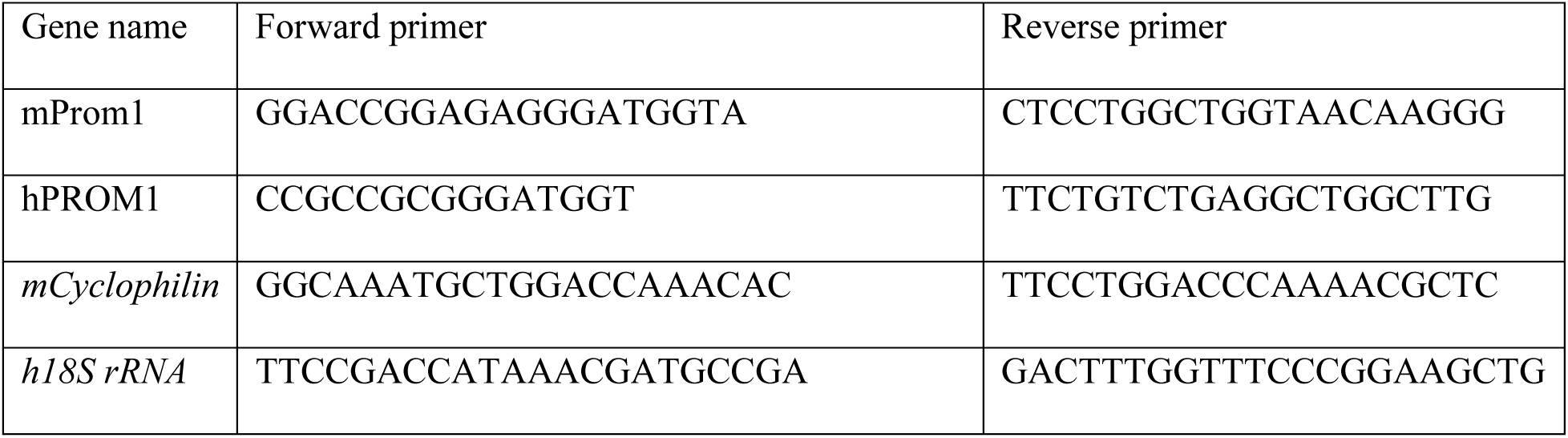
Quantitative RT-PCR primers.

## Notes

### Competing Interest Statement

The authors have declared no competing interest.

## References

1. Mocumbi A, et al. Pulmonary hypertension. Nat Rev Dis Primers. 2024;10:1.

2. Zhao H, et al. Endothelial Heterogeneity in Pulmonary Hypertension. Arterioscler Thromb Vasc Biol. 2026;46:3–16.

3. Diez M, et al. Plasticity of CD133+ cells: Role in pulmonary vascular remodeling. Cardiovasc Res. 2007;76:517–527.

4. Foris V, et al. CD133+ cells in pulmonary arterial hypertension. Eur Respir J. 2016;48:459–469.

5. Pleskac P, et al. Emerging roles of prominin-1 (CD133) in the dynamics of plasma membrane architecture and cell signaling pathways in health and disease. Cell Mol Biol Lett. 2024;29:41.

6. Peichev M, et al. Expression of VEGFR-2 and AC133 by circulating human CD34+ cells identifies a population of functional endothelial precursors. Blood. 2000;95:952–958.

7. Quirici N, et al. Differentiation and expansion of endothelial cells from human bone marrow CD133+ cells. Br J Haematol. 2001;115:186–194.

8. Salven P, et al. VEGFR-3 and CD133 identify a population of CD34+ lymphatic/vascular endothelial precursor cells. Blood. 2003;101:168–172.

9. Wang R, et al. Glioblastoma stem-like cells give rise to tumour endothelium. Nature. 2010;468:829–833.

10. Sun S, et al. CD133+ endothelial-like stem cells restore neovascularization and promote longevity in progeroid and naturally aged mice. Nat Aging. 2023;3:1401–1414.

11. Lu J, et al. Endothelial Cells Promote the Colorectal Cancer Stem Cell Phenotype through a Soluble Form of Jagged-1. Cancer Cell. 2013;23:171–185.

12. Farha S, et al. Hypoxia-inducible factors in human pulmonary arterial hypertension: a link to the intrinsic myeloid abnormalities. Blood. 2011;117:3485–3493.

13. Dai Z, et al. Prolyl-4 Hydroxylase 2 (PHD2) Deficiency in Endothelial Cells and Hematopoietic Cells Induces Obliterative Vascular Remodeling and Severe Pulmonary Arterial Hypertension in Mice and Humans Through Hypoxia-Inducible Factor-2α. Circulation. 2016;133:2447–2458.

14. Dai Z, et al. Endothelial and Smooth Muscle Cell Interaction via FoxM1 Signaling Mediates Vascular Remodeling and Pulmonary Hypertension. Am J Respir Crit Care Med. 2018;198:788–802.

15. Muzumdar MD, et al. A global double-fluorescent Cre reporter mouse. Genesis. 2007;45:593–605.

16. Buch T, et al. A Cre-inducible diphtheria toxin receptor mediates cell lineage ablation after toxin administration. Nat Methods. 2005;2:419–426.

17. Sheikh AQ, et al. Smooth muscle cell progenitors are primed to muscularize in pulmonary hypertension. Sci Transl Med. 2015;7(308):308ra159. doi:10.1126/scitranslmed.aaa9712.

18. Dierick F, et al. Resident PW1+ Progenitor Cells Participate in Vascular Remodeling During Pulmonary Arterial Hypertension. Circ Res. 2016;118(5):822–833. doi:10.1161/CIRCRESAHA.115.307035.

19. Heise RL, et al. From Here to There, Progenitor Cells and Stem Cells Are Everywhere in Lung Vascular Remodeling. Front Pediatr. 2016;4:80. doi:10.3389/fped.2016.00080.

20. Yao W, et al. Identification of putative endothelial progenitor cells (CD34+CD133+Flk-1+) in endarterectomized tissue of patients with chronic thromboembolic pulmonary hypertension. Am J Physiol Lung Cell Mol Physiol. 2009;296(6):L870–L878. doi:10.1152/ajplung.90413.2008.

21. Chettimada S, et al. Glucose-6-phosphate dehydrogenase plays a critical role in hypoxia-induced CD133+ progenitor cells self-renewal and stimulates their accumulation in the lungs of pulmonary hypertensive rats. Am J Physiol Lung Cell Mol Physiol. 2014;307(7):L545–L556. doi:10.1152/ajplung.00303.2013.

22. Solinc J, et al. Platelet-Derived Growth Factor Receptor Type α Activation Drives Pulmonary Vascular Remodeling Via Progenitor Cell Proliferation and Induces Pulmonary Hypertension. J Am Heart Assoc. 2022;11(7):e023021. doi:10.1161/JAHA.121.023021.

23. Zhang X, et al. Robust genome editing in adult vascular endothelium by nanoparticle delivery of CRISPR-Cas9 plasmid DNA. Cell Rep. 2022;38(1):110196. doi:10.1016/j.celrep.2021.110196.

24. Jin S, et al. Inference and analysis of cell-cell communication using CellChat. Nat Commun. 2021;12(1):1088. doi:10.1038/s41467-021-21246-9.

